# Aerodynamics and motor control of ultrasonic vocalizations for social communication in mice and rats

**DOI:** 10.1101/2021.03.08.434401

**Authors:** Jonas Håkansson, Weili Jiang, Qian Xue, Xudong Zheng, Ming Ding, Anurag A. Agarwal, Coen P.H. Elemans

## Abstract

Rodent ultrasonic vocalizations (USVs) are crucial to their social communication and a widely used translational tool for linking gene mutations to behavior. To maximize the causal interpretation of experimental treatments, we need to understand how neural control affects USV production. However, both the aerodynamics of USV production and its neural control remain poorly understood. Here we test three intralaryngeal whistle mechanisms - the wall and alar edge impingement, and shallow cavity tone - by combining in vitro larynx physiology and individual-based 3D airway reconstructions with fluid dynamics simulations. Our results show that in the mouse and rat larynx USVs are produced by a glottal jet impinging on the thyroid inner wall. Furthermore, we implemented an empirically based motor control model that predicts motor gesture trajectories of USV call types. Our work provides a quantitative neuromechanical framework to evaluate the contributions of brain and body in shaping USVs, and a first step in linking descending motor control to USV production.

## Introduction

Murine rodents produce ultrasonic vocalizations (USVs) that range in frequencies from 20 to 100 kHz and play a crucial role in social communication behaviors, such as mating and territorial defense (1, 2). Different USV call types strongly signal positive (3) or negative (4) emotional states (5, 6) and are crucial for pups to induce maternal search and retrieval behavior, when visual or olfactory cues are less relevant (7). USVs have been found in at least 50 rodent species (8), but are probably more widespread, given that rodents comprise over 40% of all mammal species (9) and only a fraction has been investigated (8). Furthermore, USVs have recently become an increasingly used behavioral readout in mice and rats, the two most widespread translational animal disease models in biological and medical research (10). USVs are used a translational tool for linking gene mutations to behavioral changes in rodent models for speech (11) and neuropsychiatric communication disorders, such as autism (12, 13) and Down syndrome (14). The observed changes in vocalization behavior, such as altered USV occurrence (15), sound frequency (16, 17) or aberrant USV call types (18), are attributed to changes in neural control (15–18). However, linking the brain to behavior requires a causal and quantitative understanding of the transformation from descending motor control to USV production in these species that we currently lack.

Translating motor control to USV production requires both system identification of the mechanism by which sound is produced and quantitative understanding of how muscles drive the control parameters of this system. Until recently, USVs were thought to be hole-tone whistles that require two orifices for producing a stable tone (19, 20), such as the human teeth-lip whistle and tea-kettle whistle (21). However, USVs in mice were recently shown to be produced by a sound production mechanism novel to mammals and previously only identified in industrial supersonic and high-speed subsonic flows (22–24): a glottal jet impinging on a structure within the larynx (25). Small instabilities in a glottal air jet that travel downstream are entrained to occur at certain frequencies due to a feedback loop between these downstream-travelling flow structures and acoustic waves travelling upstream. In the small murine larynx, where glottal jet speeds can reach up to 10% of the speed of sound, the jet impingement mechanism can lead to stable high frequency tones from 20 to over 100 kHz (20, 25, 26). The impingement structure within the larynx has been proposed to be either the thyroid wall (25) or a laryngeal adaptation (27) found in several muroid rodents, the alar edge (27, 28) (Fig. 1). Both mechanisms constrain motor control to the respiratory and laryngeal musculature, but the proposed aeroacoustic models for wall and alar edge tones occur under distinct physiological conditions and predict very different sound frequencies (25, 27). Thus establishing which aerodynamic mechanism is responsible for USV production is critical for quantitatively linking neuromuscular control parameters to USV acoustics.

**Figure 1.**
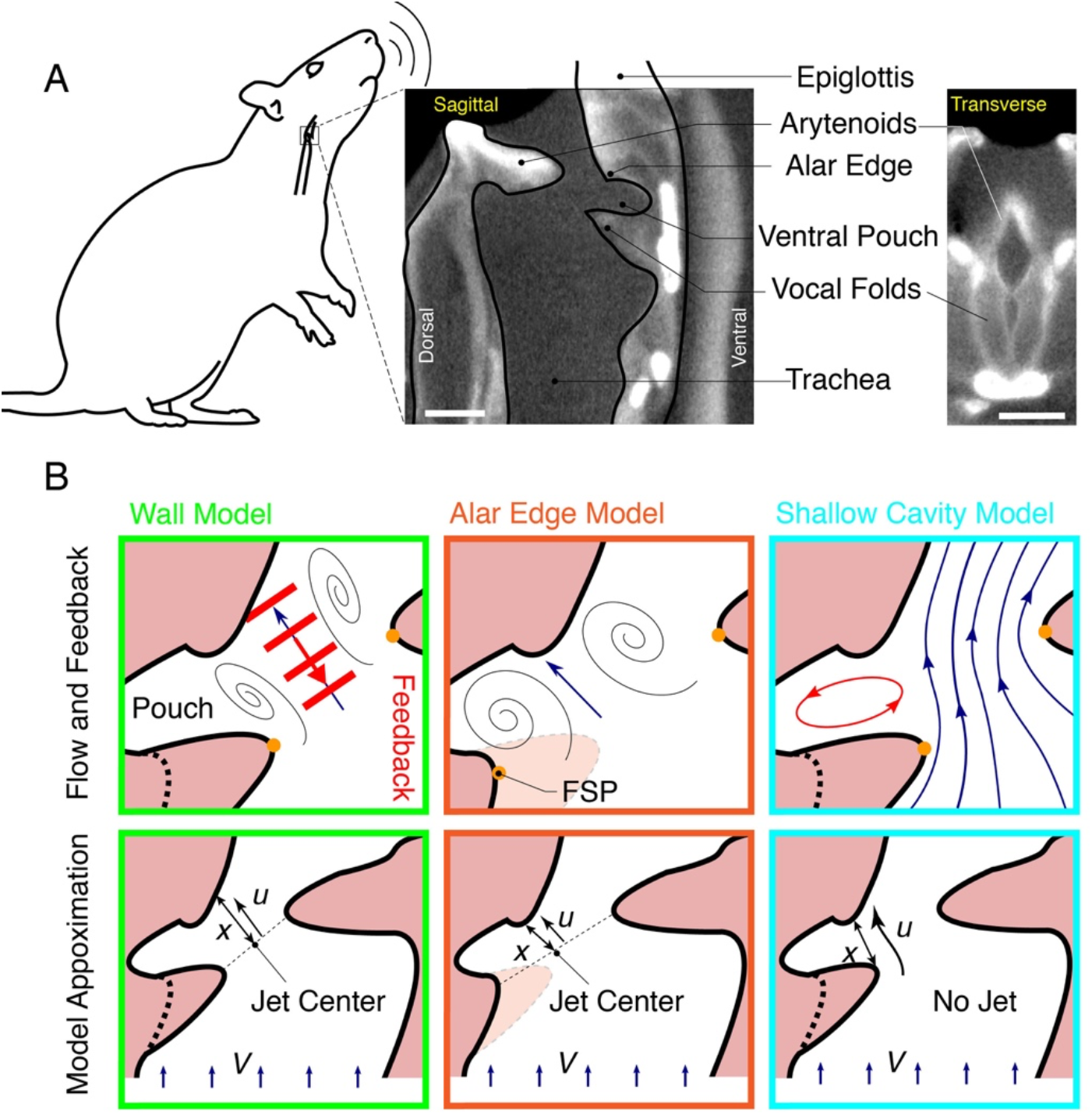
Proposed aeroacoustic mechanisms of USV production in the rat and mouse larynx. (A) Dice microCT scan of the rat larynx with cross-sections in medial sagittal plane (middle), and transversal plane parallel to the vocal folds (right). (B) Schematic of wall impingent (left), alar edge (middle), and shallow cavity (right) aerodynamic mechanisms of USV production in rats. The models are distinct in their local flow conditions (top row, black lines), feedback mechanism (red) and model parameters (bottom row) with jet impingement length x, jet exit speed u, and tracheal flow V. FSP, Flow Separation Point (orange dots).

Here we test what aerodynamic mechanism explains USV production in rats and mice. We exploit different predictions made by the main two proposed mechanisms – wall and alar edge impingement tones – and furthermore introduce a novel mechanism, the shallow cavity tone, that we propose as a more likely aerodynamic flow scenario than an alar edge tone. We combine a series of *in vitro* excised larynx experiments with computational flow models to test three distinguishing key physiological boundary conditions. We show that USVs are produced with adducted vocal folds, that only the wall impingement model predicts anatomically correct glottal air jet parameters, and that normal USVs are produced in absence of the alar edge and ventral pouch. Together, all datasets strongly support the intralaryngeal wall impingement mechanism. We then propose a quantitative motor control model that derives time-resolved control parameters from *in vivo* USV sound recordings and provides a physiological basis for USV syllable categorization and interpreting rat and mouse vocal behavior phenotypes. Our model furthermore shows that both brain and body contribute to USV frequency traces which emphasizes the importance of an embodied or systems approach to USV motor control.

## Results and Discussion

We tested three physiological boundary conditions that are distinctive between wall and alar edge tone models for USV production: *i)* vocal fold adduction state, *ii)* jet separation and impingement locations, and *iii)* the presence of the alar edge and ventral pouch cavity.

The first distinctive feature between wall and alar edge tones is vocal fold adduction state (Fig. 1B). In mice, USVs are produced *in vitro* with fully adducted (opposed) vocal folds, which leaves a glottal opening on the dorsal side between the arytenoid cartilages, i.e. the cartilaginous glottis, for respiratory flow to go through (25). In contrast, the alar edge tone model predicts tones to occur with vocal folds abducted (open), resulting in a much larger glottal opening that includes the ventral opening between the vocal folds (i.e. the membranous glottis) plus the cartilaginous glottis (27).

To test which vocal fold adduction state leads to USV production in rats, we used an excised larynx paradigm that allowed detailed manipulation of glottal configuration (25, 29) (*Methods and Materials, Additional File 1*). We subjected rat larynges to pressure ramps with abducted and adducted vocal folds. With adducted vocal folds all larynges produced fictive USVs (fUSVs) (N = 10), while only 1 out of 10 produced fUSVs with abducted vocal folds. fUSVs were produced over a phonation threshold pressure of 0.8±0.3 kPa (N=10), consistent with *in vivo* values of 0.4-0.9 kPa (30). Flow ranged from 2.6±0.6 – 3.7±1.0 ml/s (N=10), which is within estimated physiological range of 0-10 ml/s (*Methods and Materials*). Furthermore, the fUSVs peak frequencies ranged from 25-61 kHz, which corresponds well to the *in vivo* range of 18-96 kHz, including “22 kHz” (range 18-32 kHz) and “50 kHz” (range: 32 - 96 kHz) USVs (31). Thus, driving excised rat larynges with physiologically realistic airflows cause fUSVs that overlap in acoustic parameters with *in vivo* USVs, which suggest that the *in vitro* paradigm represents the *in vivo* situation very well. Our data supports the hypothesis that USVs in rats are produced with adducted vocal folds, which is consistent with a reduced airflow during USV production compared to quiet respiration in rats (30, 32), earlier *in vitro* glottal adduction manipulations that lacked sound recordings (33) and preliminary *in vivo* endoscopic observations (34). Thus, USVs in both mice (25) and rats are produced with adducted vocal fold, which provides evidence against the alar edge tone mechanism and in favor of the wall tone mechanism.

The second distinctive feature between wall and alar edge models is the speed, position, length, and angle of a formed air jet. The wall tone model predicts jet formation at the center of the cartilaginous glottis and impingement on the thyroid inner wall (Fig. 1B) (25). The alar edge tone model predicts that “the glottal jet exits close to the ventral side of the laryngeal lumen, resulting in a glottal jet path nearly parallel to the intralaryngeal supraglottal wall” (27). Thus, the required jet is proposed to separate on the ventral side of the laryngeal lumen (at Flow Separation Point, FSP, Fig. 1B), which implies that the jet center is located at the center of the glottis (Fig. 1B). Jet impingement is constrained to the alar-edge (27). These jet location differences thus result in different jet angles and lengths, which in turn lead to different flow-frequency transformations (*Methods and Materials*). However, we think the proposed alar edge model poses an unlikely flow scenario for the formation of a separated jet - essential to the edge tone - because the large glottis leads to low flow speeds and a low flow constriction ratio. We also question the validity of the assumption that the pouch can act as a Helmholtz resonator (27), because the anatomical structure to act as the essential neck is not present. Instead, we propose a third USV production mechanism, the shallow cavity tone, which is based on a more realistic flow scenario that does not require jet formation, has FSP at the same location as the alar edge model, and leads to stable high-frequency whistles (35). Cavity flows are produced when air flow detaches flows over a cavity and reattaches downstream of the cavity (at the thyroid in Fig. 1B) and sets up a recirculating flow inside the cavity. The flow can produce loud tonal sounds. Such flows are of significant interest in aerospace applications, such as wheel wells and weapon bays of aircraft, where the strong oscillations from the tones can lead to significant structural damage (36).

To estimate flow and jet conditions, we combined USV production under controlled *in vitro* conditions with morphometric analysis of individual-based dice-CT scans. In all models, frequency is set by *u*, the mean convection speed of the coherent flow structures, approximated as the glottal exit speed, and jet or cavity length (*Methods and Materials*). While the cavity tone model does not predict the formation of a jet, it does rely on the length of the entrance to the ventral pouch and thereby, for a given frequency, also predicts a length. We measured tracheal airflow (*V*) and peak frequency (*f*_*p*_) during fUSV production (Fig. 3) in fresh larynges (N=5) that were subsequently fixed in PFA to stabilize the geometry. Even after PFA fixation, fUSVs were produced in all larynges and the slope of the frequency-to-flow relationship did not differ significantly before and after fixation (Paired t-test, 2-tailed, p = 0.48). We subsequently measured the glottal area (*A*_*gl*_) in Dice-µCT scans (Fig. 3A, *Methods and Materials*) of all individuals to estimate jet exit speed *u*. The produced frequencies and jet speed predicted jet lengths of 0.92 ± 0.21 mm, 0.46 ± 0.11 mm, and 0.46 ± 0.13 mm for the wall, alar edge, and cavity tones respectively (Fig. 3C), and jet angles of 99.2 ± 15.3° and 62.3 ± 11.1° (n=5) for wall- and alar edge-tone, respectively (Fig. 3D). Jet angle was not predicted by the cavity tone model. To test if these predicted lengths were consistent with the physical dimensions of the larynx, we measured minimum wall jet length (x_*wall*_), alar edge jet length (x_*alar*_), and cavity length (x_*cav*_) on Dice-µCT scans of the individual larynges (Fig. 3A). For the wall impingement model the predicted jet lengths were not significantly different from the minimum length (two-tailed paired sample t-test, p = 0.09), and importantly fell within the physical range in all five cases (Fig. 3C). However, the predicted jet length for the alar edge-tone model was significantly shorter than the anatomical length (two-tailed paired sample t-test, p = 0.003), and fell 0.26 ± 0.07 mm too short to reach the alar edge (Fig. 3C). The predicted cavity length for the cavity-tone model was also significantly shorter than the anatomical length (two-tailed paired sample t-test, p = 0.020), (Fig. 3C). Therefore, these experiments support the wall-tone whistle mechanism.

**Figure 2.**
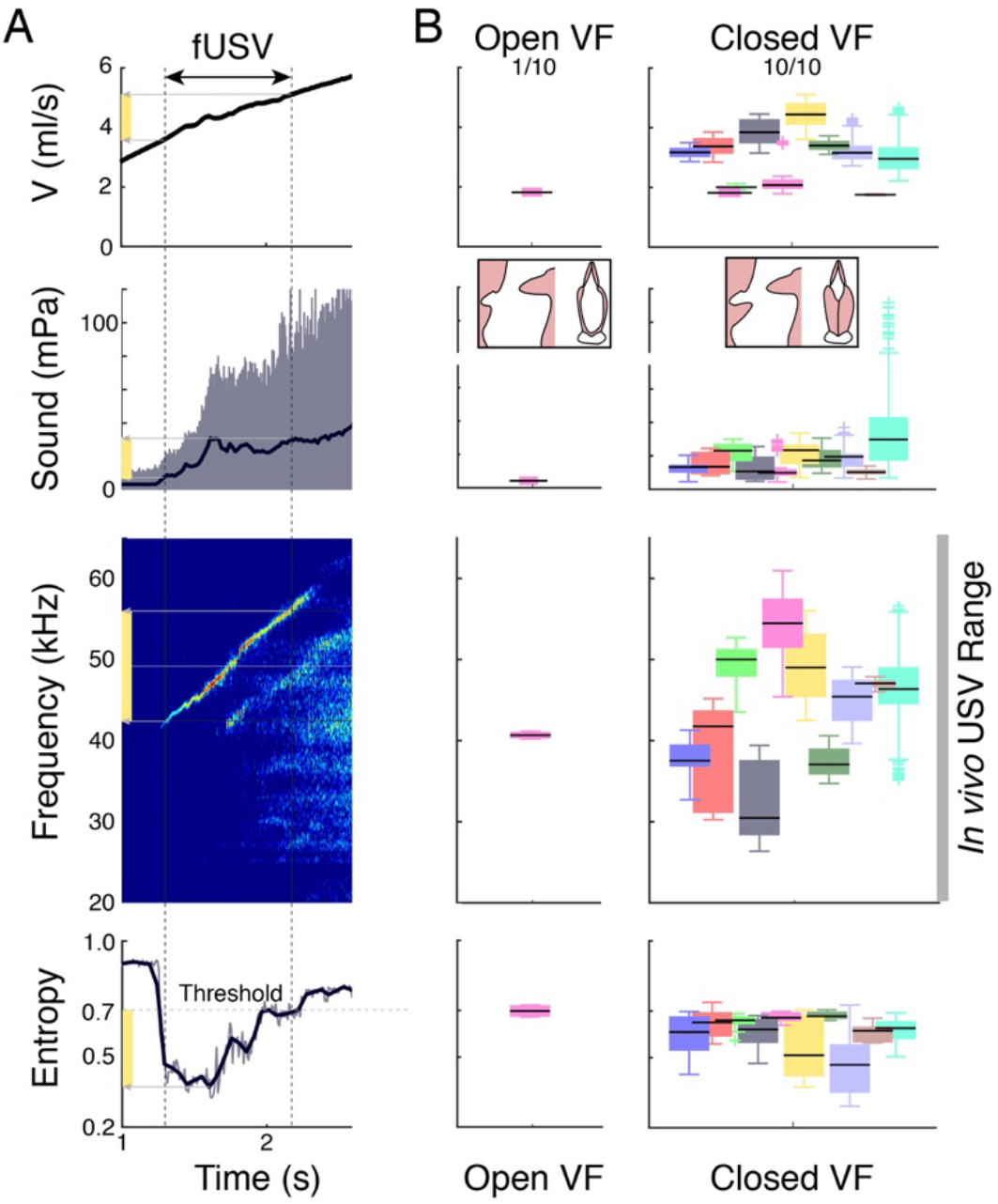
Rat fUSVs are produced with adducted vocal folds. (A) Above a threshold tracheal flow *V*, the isolated larynx produces fUSVs. From top to bottom: tracheal mass flow *V*, received sound pressure, sound spectrogram (NFFT=2048, overlap=50%, Hamming window), and scaled Shannon’s entropy with the 0.7 threshold for USV detection. (B) With abducted vocal folds and open membranous glottis only 1 larynx produced fUSVs (left), while with adducted, opposed vocal folds all larynges (N=10) produced USVs (right) and within the *in vivo* frequency range of 18-96 kHz (31). Different colors represent different individuals.

**Figure 3.**
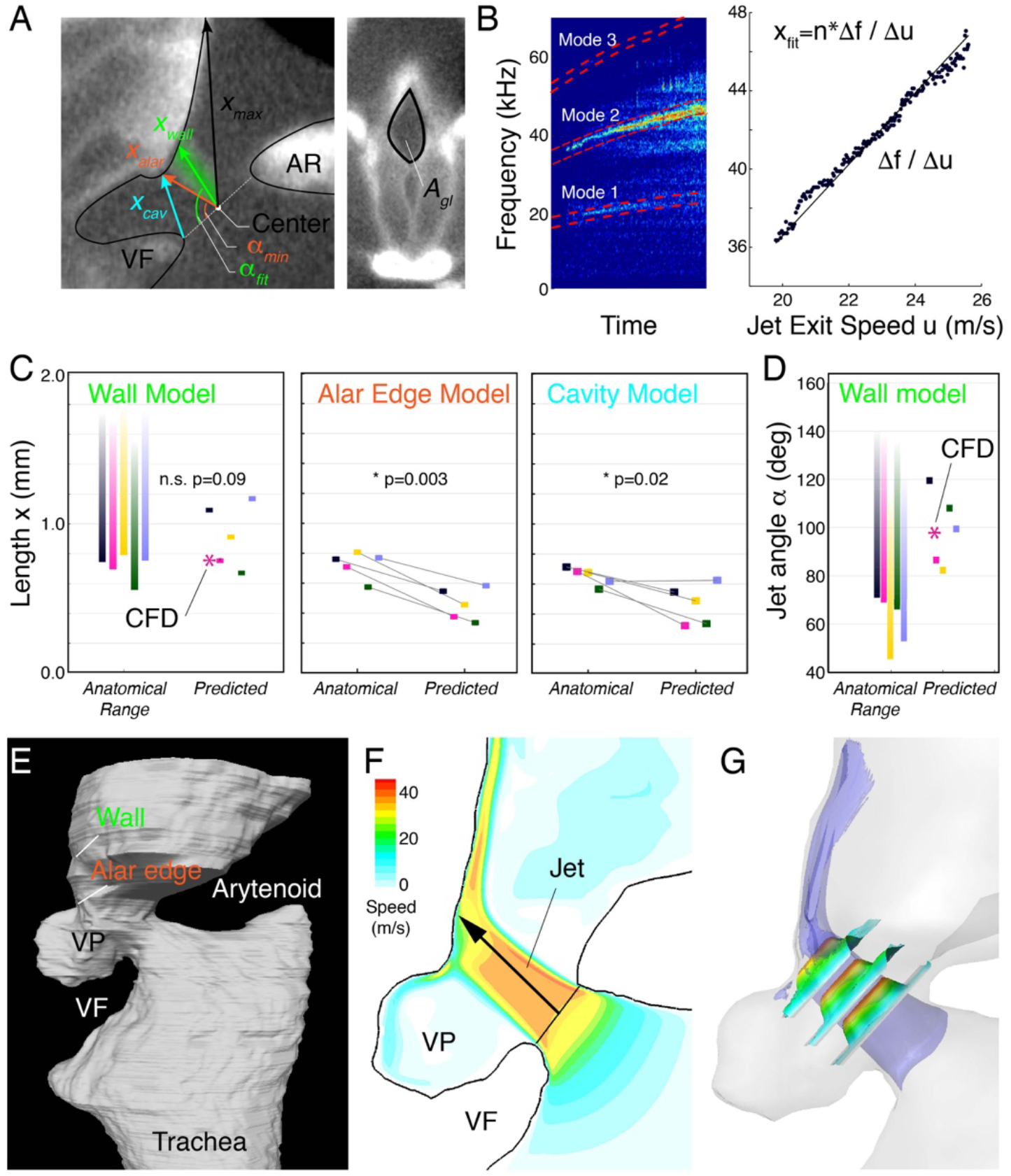
Glottal jet parameters support wall impingement model in rats. (A) The anatomical lengths of wall (*x*_*wall*_) and alar edge (*x*_*alar*_) jets, and ventral pouch cavity opening (*x*_*cav*_) as measured in sagittal cross-sections of the glottis (left). Area of the cartilaginous glottis (*A*_*gl*_) was measured in a transverse section parallel with the glottal opening (right). (B) Spectrogram (NFFT=2048, overlap=50%, Hamming window) of a fUSV shows multiple modes (red dashed boxes) essential to determine the dominant mode (See *Methods and Materials*). The slope between dominant frequency and jet speed equals the predicted jet/cavity length *x* (right). (C) Observed anatomical versus predicted values for *x* in wall, alar edge, and cavity model and (D) jet angle. These data show that wall-tone jet length and angle predictions fall within, while alar edge and cavity model predictions fall below the anatomical length range (C) or do not provide a solution for angle (D). (E) Flow was simulated in a fixed 3D mesh of the laryngeal airway. (F) 2D and (G) 3D flow show that a distinct jet is formed and impinges on the thyroid wall. Blue; iso-surface of jet speed equals 30 m/s. The three small planes present speed profiles and are contoured also by the speed value.

To further test if the predicted jet length and angles were consistent with intralaryngeal flow, we performed Computational Fluid Dynamics simulations (37) of airflow through a 3D-reconstructed larynx in USV producing state (Fig. 3E, See Methods and Materials). Using the same boundary conditions as under experimental settings, our CFD model showed first of all that jet formation occurred with jet separation points at the dorsal and ventral side of the cartilaginous glottis (Fig. 3F,G; Additional File 2). Second, the jet impinged on the thyroid planar wall and not the alar edge. Third, the jet was 0.76 mm long, at a 98.0° angle and had a speed of 36.5 m/s, which was in excellent agreement with the predicted *x*_*wall*_ = 0.71 mm at 86.6° and 33.2 m/s of our aeroacoustic model for that individual (Fig. 3E-G). The simulated jet angle is also in excellent agreement with the earlier estimate in the mouse larynx (25). Taken together, the predicted jet lengths and flow structure from CFD simulation provide evidence against both alar-edge and shallow cavity-tone models, and support the intra-laryngeal planar impinging jet model of USV sound production in rats.

The third distinct feature between the wall, alar edge and cavity tone models is the required presence of the alar edge and a small airsac-like cavity rostral to the vocal folds, called the ventral pouch, which is found in several muroid rodent species (27, 38, 39). The wall tone model allows air circulation in the ventral pouch, but does not require its presence because the feedback that stabilizes the tone comes from acoustic waves within the jet (22–25). The alar edge tone model on the other hand evidently requires the presence of the alar edge and suggests that pouch cavity resonance properties affect sound frequencies (27). The cavity tone model too requires the presence of the ventral pouch for air circulation. To test if the alar edge and ventral pouch are essential to fUSV production, we prevented both the presence of an edge and air circulation in the pouch by filling the pouch with a small aluminum sphere in excised rat (n=7) and mice (n=6) larynges. In rats, six out of seven larynges retained fUSV production after sphere insertion (Fig. 4) and the mean, minimum, and maximum peak frequencies (*f*_p_) did not change significantly in rats (two-tailed paired sample t-test, N=7, mean *f*_p_: p = 0.89, max *f*_p_: p = 0.71, min *F*_p_: p = 0.87) and mice (N=6, mean *f*_p_: p = 0.65, max *f*_p_: p = 0.48, min *f*_p_: p = 0.45). To estimate how filling the ventral pouch affected the intralaryngeal flow, we performed CFD simulations of the same experimental manipulation (Fig. 4E,F; Additional File 3). A glottal jet formed that impinged on the thyroid planar wall slightly more rostral due to the sphere, leading to a slightly increased angle (103°, +5.1%) and jet length (0.79 mm, +2.6%). Thus, neither the ventral pouch or the alar edge is essential for USV production in rats and mice

Finally, we used CFD simulations to test if the proposed flow scenario (27) for the alar edge model in Fig 1B is physically plausible. We ran CFD simulations on the previously published 3D reconstructed rat vocal tract (27) that has abducted VFs and arytenoids (See methods). Driven by *in vivo* tracheal pressure, our simulations show that no intralaryngeal jet is formed and no air circulates in the ventral pouch (Additional File 4). Therefore we can conclude that the suggested flow scenario for the alar edge model (27) is not physically accurate.

**Figure 4.**
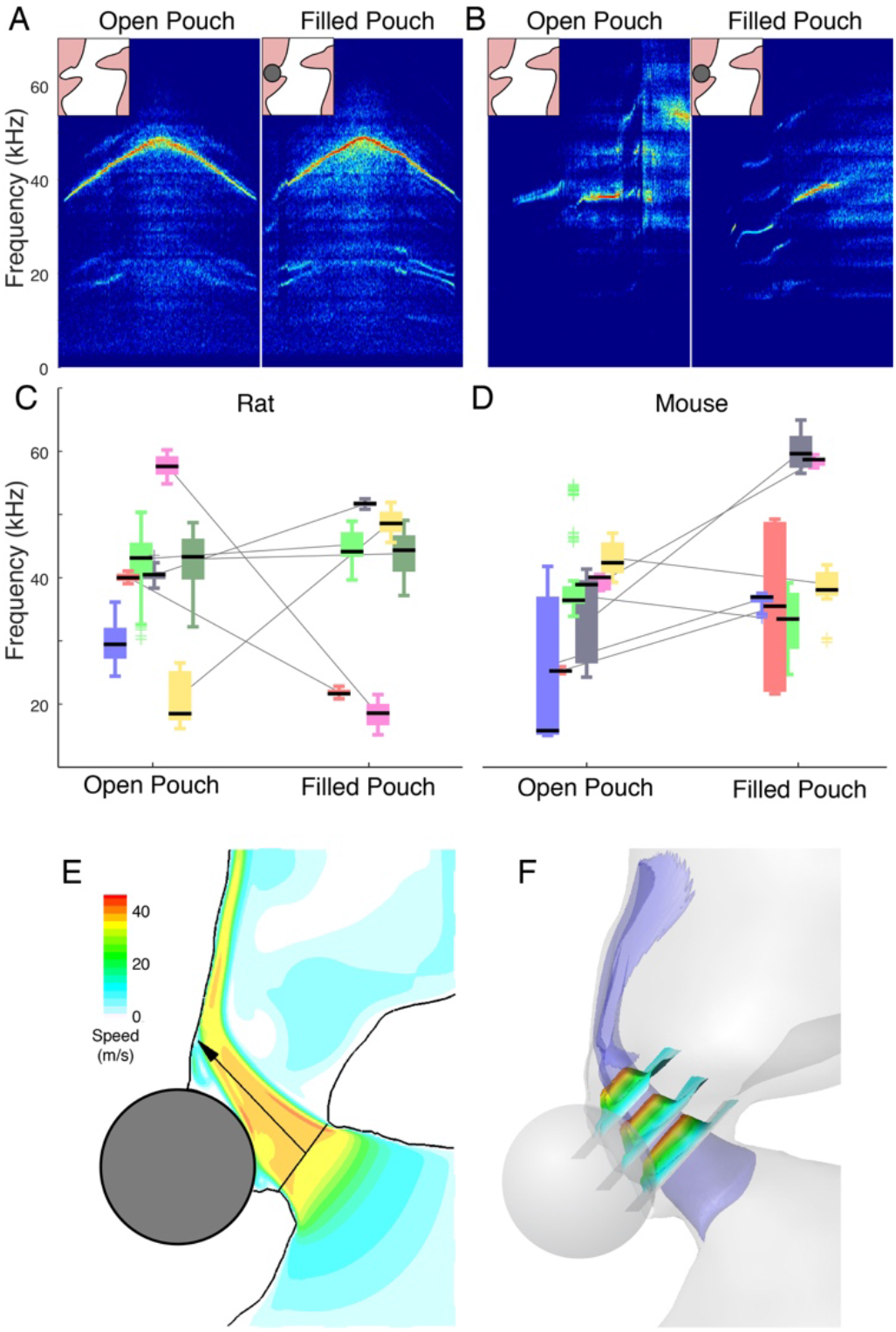
The alar edge and ventral pouch are not required for USV production in rat and mouse larynx. Example spectrograms of normal fUSVs (left) and blocked alar edge and filled ventral pouch (right) by small aluminum sphere in (A) rat and (B) mouse larynx. (C) Six out of 7 rat larynges and (D) 6 out of 6 mice larynges produced fUSVs with filled ventral pouch. (E) Computational fluid dynamic simulations in the rat larynx slice and (F) 3D rendering show that also with a filled ventral pouch, a jet forms that impinges on the thyroid wall with negligible effect on the jet length and angle.

Our data conclusively shows that both in the most widely used rodent models in biological and medical research, rats and mice, USVs are produced by an aerodynamic wall impingement whistle. The three distinctive features - closed vocal fold adduction state, jet properties and non-essential presence of edge and pouch - provide evidence against alar edge and shallow cavity tones and support the wall tone. The notion that wall impingement is incongruent with laryngeal anatomy (27) is thus incorrect. However, given the large diversity of laryngeal morphology and life history found in the 1500 species of rodents (40), our data does not exclude that multiple mechanisms contribute to USV production in other rodents species such as singing (41, 42) or grasshopper mice (43). Shallow cavity tones (35) provide an alternative mechanism to explain the loud USVs of several new world rodents with more pronounced alar and pouch structures and may be a wide-spread mammalian sound production mechanism that requires further investigation.

In vivo rodent USVs are characterized and classified by the time-varying frequency trajectories of syllables (18, 30 31). Based on our aerodynamic model of USV production, we have implemented a quantitative data-driven model of in vivo USV motor control (Methods and Materials). Our aerodynamical model of USV production predicts that the frequency of pressure and flow structure variations are set by the jet speed and jet length. The frequency of these whistles is 50-100 kHz and the pressure fluctuation thus occur at the microsecond scale and are at least three orders of magnitude faster than the millisecond laryngeal motor control (30, 44, 45) of the jet parameters jet speed and jet length. As a consequence, the USV instantaneous frequency can be considered time-invariant compared to the motor control that shapes the frequency trajectories. Furthermore, in contrast to an earlier suggestion (27), the fact that USV exhibit changes *in vivo* does thus not inform on the aerodynamical mechanism. We focused on rats where pressure, flow and muscle electromyography data has been measured during USV production *in vivo* (30, 44, 45). Within correct anatomical and physiological ranges, the *x, u* control space produces all frequencies observed *in vivo* (Fig. 5A,B). We used an orifice constriction model that accurately estimated tracheal mass flow from pressure (Fig. 5C) to calculate how subglottal pressure and glottal area affect frequency (Fig. 5D). Surprisingly, glottal area barely affects frequency because the increase in jet speed from decreasing area is counteracted by the decrease in flow.

**Figure 5.**
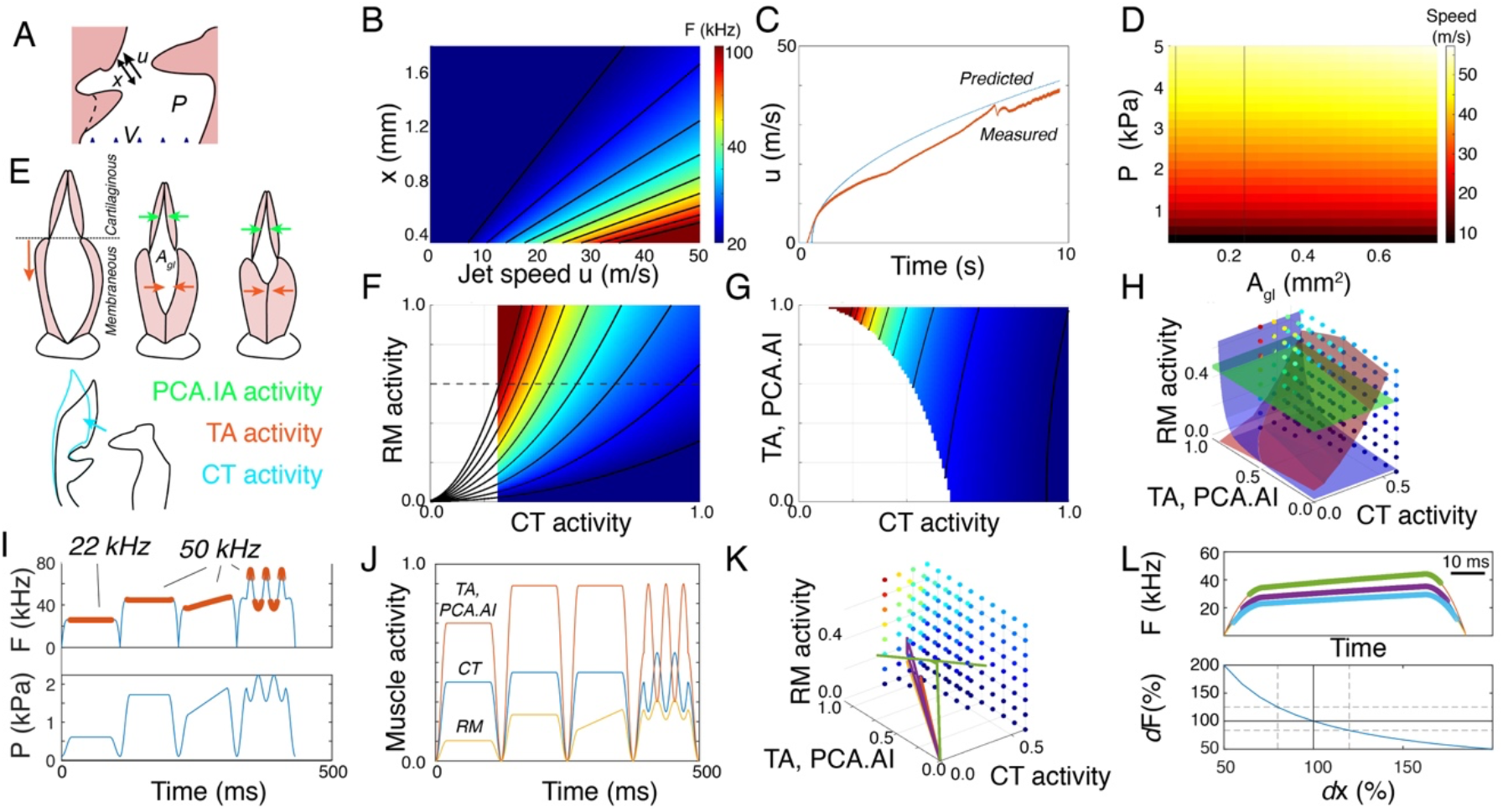
Embodied motor control model extracts motor gestures for rat USVs call types. (A) Exploring the parameter space of our aerodynamic wall impingement model results in (B) frequencies that overlap with in vivo range. Impingement length (x) and jet speed (u), tracheal flow V. (C) Predicted flow by orifice obstruction model (blue) corresponds well to measured flow (red) during subglottal pressure ramp through rat larynx in vitro. (D) Jet speed (u) as function of glottal area shows little dependency on glottal area in relevant range (vertical black lines represent 0.5 and 2 times glottal areas from CT scans), whereas subglottal pressure strongly influences jet speed. (E) Effects of muscle shortening on laryngeal geometry (See *Methods and Materials*). Top, combinations of intrinsic laryngeal muscles affect the membranous and cartilaginous glottal area. Bottom, *m. cricothyroid* (CT) contraction leads to thyroid rotation (black to cyan outline), which increases impingement length. The rotatory action of CT is assumed to weakly counteracted by the smaller thyroarytenoid (TA) muscle. (F) Both respiratory muscle (RM) and CT activity affect USV frequency. The whistle is unstable in the white area. Black horizontal dashed line indicates the upper subglottal pressure limit during USVs in vivo (*p*_*sub*_ = 3 kPa). (G) TA action strongly influences the stability of the whistle and as such gates sounds, while it has little effect on *f*_*0*_. CT action affects both stability and *f*_*0*_. (H) Frequency *f*_*0*_ is highly redundant in the three-dimensional motor space (red isosurface; *f*_*0*_ = 45 kHz). Adding a given subglottal pressure (green isosurface; *p*_*t*_ = 2.5 kPa) and flow (blue isosurface; *V* = 4.2 ml/s) reduces this redundancy into a line or single point. (I) Driven by USV frequency (top; orange stable frequencies) and subglottal pressure (bottom), our model predicts muscle activity in (J) time and (K) as gestures in motor space for two common USV call types, 22 and 50 kHz, including the subtypes with frequency modulations and jumps. (L) Small changes in larynx geometry, such as impingement length *x*, alter the contours (top) and frequencies (bottom) of USVs whilst driven by identical motor gestures.

Two motor systems drive the parameters of our model; first, the respiratory muscles that control subglottal pressure and second, intrinsic laryngeal muscles that control laryngeal geometry, such as glottal area and impingement length. Because rodent laryngeal muscles share developmental origin (46), location and function (39) with other vertebrates, we based their mechanical actions on better studied mammals such as human (47) and dog (48–50). We included three muscle groups; respiratory muscles (RM) that control subglottal pressure, the cricothyroid muscle (CT) that controls impingent length, and a combination of intrinsic laryngeal muscles (Thyroarytenoid (TA), posterior cricoarytenoid (PCA) and interarytenoid (IA) muscles) that set vocal fold adduction and thereby glottal area (Fig. 5E; *Methods and Materials*). Consistent with earlier observations in mice (25), with increasing CT force and decreasing *x*, USV frequency goes down. Interestingly, the CT has thus the opposite function compared to vocal fold vibration driven voiced sound production where CT shortening increases frequency (29, 49, 51). The laryngeal muscles affect the jet shape and flow conditions that determine whistle stability, which gates the sound on and off (*Methods and Materials*). The three muscle groups together affect USV frequency in a highly redundant control space (Fig. 5F-H), which makes it challenging to invert the system and estimate control parameters from sound alone. However, with additionally known factors such as pressure or flow, and at higher frequencies where the jet becomes unstable, this redundancy collapses (Fig. 5H).

We computed putative *in vivo* motor control trajectories of 22 kHz and 50kHz USV calls (6) from acoustics and corresponding *in vivo* subglottal pressure (30, 44) (Fig. 5I-K). Our model can reproduce these call types including several subtypes, such as flat and modulated trill calls including frequency jumps (Fig. 5I), and accurately predicts that pressure increases and flow decreases during USVs consistent with *in vivo* recordings (30, 32, 52). Moreover, increased TA and CT force correlates with higher frequencies (Fig. 5K) consistent with *in vivo* recordings (44) to counteract abductive forces of increased respiratory pressure and to overcome whistle instability. Lastly, we explored the effects of changing larynx geometry with unchanged motor control trajectories and show that change of only 180 µm (20%) can cause frequency shifts of 10 kHz, which is similar magnitude observed in behavioral models (16, 17). Taken together our simple model provides a physiological basis for the neuromuscular control of USVs and interpreting rat and mouse USV call phenotypes.

Mice and rat USVs often contain distinct frequency jumps that play an important role in call type classification (2, 53). These jumps occur on the millisecond scale and do not correlate with either laryngeal muscle activity or pressure (30, 45). Our aerodynamic model predicts that these frequency jumps are jumps between stable whistling modes which explains why they can overlap *in vivo* (53). Our motor model includes jet stability criteria that predict which simultaneous modes are stable, and these seem to correspond well (Fig. 5I) with *in vivo* observed jumps in rats (2). What exact modes are finally produced *in vivo* depends on local flow conditions at the jet exit and needs further investigation.

Detailed control of laryngeal muscles is crucial in shaping USVs and connecting spiking motor neurons to muscle action and laryngeal biomechanics requires more complex modelling approaches and additional knowledge on motor neuron and muscle properties, motor unit organization, and mechanical effects of muscle shortening. Interestingly, laryngeal muscles are typically very fast, but the cricothyroid muscle’s origin is somatic and it is slower than e.g. TA in many mammals including rats (54, 55). Because our model suggest that USV frequency is predominant set by CT action and respiratory pressure, both rather slow systems compared to the precision and speed of other vocal production system, such as birdsong (56, 57) and bat echolocation (58), this may explain the slower cadence of frequency modulation in in rat and mice USVs.

The brain is constrained and modulated by the biomechanics, morphology and material properties of the body (59–61). Our findings show that both neural and anatomical components contribute to USV production (Fig. 5L). Therefore, the mechanisms that drive changes in strain specific USVs or USV changes in mouse and rat disease models can be both altered motor programs and laryngeal geometry. This emphasizes the importance of an embodied approach to USV motor control to provide a physiological basis for USV syllable categorization and interpreting rat and mouse vocal behavior phenotypes.

## Materials and Methods

### Subjects

We used 16 male Sprague Dawley rats, 11 juveniles (51 to 78 days old) and 5 adults, and 6 adult male C57BL/6 mice. All animals were housed at Odense University Hospital. All experiments were conducted at the University of Southern Denmark and were in accordance with the Danish Animal Experiments Inspectorate (Copenhagen, Denmark).

### Larynx dissection and mounting

All animals were euthanized with fentanyl/fluanisone or carbon dioxide, and kept on ice (maximally 180 min). The trachea, larynx, and surrounding tissue were dissected, flash frozen in liquid nitrogen and stored at -80°C. Before each experiment, we thawed the tissue in a refrigerator and then submerged it in refrigerated ringer’s solution (62) in a dish on ice and removed additional tissue surrounding the larynx and trachea. We then mounted the larynx in the setup. For rats, we mounted the larynx on a plastic Luer connector (1.1mm inner diameter and 1.6 mm outer diameter), filed down so that the tip was a straight tube. For mice, we mounted the larynx on a rounded, blunt 19G needle. The larynx was slid over the tube connector until the caudal edge of the cricoid touched the tube exit and secured with a suture around the trachea, 6-0 braided silk suture (Deknatel, USA) for rats, and 10-0 monofilament suture for mice.

### Experimental setup

We mounted larynges in an excised larynx setup described in detail in (25). In brief, this setup (Figure S1), allows for running humidified air through the larynx at precisely controlled pressure, while simultaneously measuring volumetric flow, pressure, and sound. The position of arytenoid flanges is controlled by micromanipulators. The rate of volumetric flow through the larynx was measured using a MEMS flow sensor (PMF2103, Posifa Microsystems, San Jose, USA). Sound was recorded using a 1/4-inch pressure microphone-pre-amplifier assembly (model 46BD, frequency response ±1dB 10 Hz – 25 kHz & ±2dB 4 Hz – 70 kHz G.R.A.S., Denmark) located 5 cm above the larynx pointing downwards and to the side of the larynx as not to be hit by the airflow leaving it (Fig. S1). The microphone signal was amplified by 40 dB for rats and 50 dB for mice (amplifier 12AQ, G.R.A.S., Denmark). We calibrated the microphone before each experiment (Calibrator 42AB, G.R.A.S., Denmark). The positions of the larynx and microphone were fixed relative to each other (Fig. S1). The sound, pressure and flow signals were low-pass filtered at 100, 10 and 10 kHz, respectively (filter model EF502 low pass filter DC – 100 kHz and EF120 low pass filter DC – 10 kHz, Thorlabs, U.S.A.) and digitized at 166, 224 (mice), or 240 kHz (USB 6259, 16 bit, National Instruments, Austin, Texas). All control and analysis software were written in MATLAB 2018a (MathWorks).

We imaged laryngeal configuration during ramps with a Leica DC425 camera mounted on a stereomicroscope (M165-FC, Leica Microsystems) or with a high-speed camera (MotionPro X4-M-4, Integrated Design tools, Inc., USA) at 250 fps. The DC425 camera was controlled using LAS (Leica Application Suite Version 4.7.0, Leica Microsystems, Switzerland), and the high-speed camera was controlled using Motion Studio (x64, Version 2.10.01, Integrated Design tools, Inc., USA). We illuminated he larynges with Leica GLS150 lamp (photography) or Leica EL6000 (highspeed imaging) through a liquid light guide connected to the stereomicroscope.

### Experimental protocols

We performed three experiments to study USV production in the larynx *in vitro*. In all experiments, we applied air pressure ramps from 0 to up to 2 kPa.

#### Protocol 1 – Vocal fold adduction

We first applied a pressure ramp in resting state without any vocal fold or arytenoid adduction. Because the airflow typically pushed the arytenoid flanges apart, we next approximated the arytenoid flanges with suture (Suture: Black Polymaide Monofilament USP. 10-0 (0.2 metric) 13 cm, Needle: Taper Point, 4mm, 70 *µ*, 90°. AROSurgical Instruments Corporation, USA) to stabilize the glottis dorsally. Next, we applied pressure ramps with 1) the vocal folds in rest position and 2) with the vocal folds adducted using two adduction methods. First, we adducted the vocal folds using micromanipulators. Next we glued the vocal folds in adducted state by applying cyanoacrylate tissue glue (3M Vetbond, TissueAdhesive – 1469-SB, 3M Animal Care, U.S.A) with a pulled glass micropipette to the rostral side of the vocal folds in an adducted state. We recorded the glottal configuration using high-speed video (250 fps) and still image camera for 6 and 4 larynges, respectively. We obtained complete datasets in 10 rats.

#### Protocol 2 – USV production in fixed larynges

After the last ramp of Protocol 1, for five animals we coated the outside of the larynx in UV-glue (Loon outdoors, UV FLY clear finish, thick, USA) and placed the larynx and mounting tube in 4% PFA. After 7 days, we mounted the fixed larynx in the setup and applied a pressure ramp to test if fUSVs were produced.

The larynx was stained for two days in 15% Lugol solution, 1 day in 10% Lugol solution, and 1 day in 5% Lugol solution on a roller mixer (Stuart SRT6D, Cole-Parmer, UK) at 6 rpm. The samples were then rinsed in distilled water for 2 times 10 minutes on the roller mixer at 12 rpm, and scanned in a µCT scanner (µCT50, Scanco Medical AG, Brüttisellen, Switzerland, 8 µm resolution) at Odense University Hospital. We obtained complete datasets in 5 rats.

#### Protocol 3 – USV production with filled ventral pouch in rats and mice

We applied pressure ramps with subsequently 1) the vocal folds and arytenoids adducted (as in Protocol 1), and 2) with an aluminum sphere placed in the ventral pouch. This size sphere fitted exactly in the pouch to fill it completely and caused the alar edge to lay on top of the sphere (Fig. 3B).

Based on measurements from CT scans, we used a 0.8 mm diameter sphere for rats, and a 0.5 mm sphere for mice, to fill the pouch. We then subjected the larynges to a pressure ramp. For rats, we used a ramp from 0 to up to 1.5 kPa and down to 0 kPa again at a rate of 0.5 or 1.66 kPa/s. For mice, we used a ramp from 0-2 kPa at a rate of 0.5 kPa/s. The position of the sphere in the ventral pouch was confirmed with a photo before and after the pressure ramp We obtained complete datasets for 7 rats and 6 mice.

### Signal conditioning and USV extraction

Digitized sound signals were bandpass filtered (3^rd^ order Butterworth filter; 2.5-100 kHz (Protocol 1, 2, and 4 rats and all mice in protocol 3) or 2.5 - 83.3 kHz (3 rats in Protocol 3) with zero-phase shift implementation *filtfilt* function). We calculated spectrograms (nfft = 2048 or 1024 (for 3 rats in Protocol 3), overlap: 0%, Hamming window). For each time bin, we calculated mean flow (*V*) and Shannon’s entropy (63) scaled to *log2(nfft2/2)* of the spectrogram’s power distribution between 15-100 kHz (All but 3 rat individuals in protocol 3) or 15-82.3 kHz (3 rat individuals in Protocol 3). Entropy was averaged over six time bins for rats, and three time bins for mice. Because turbulent air flow at high flow rates produces white noise up to 100 kHz, we designed a subjective detector for USV whistles over flow-induced noise. We used the pressure ramps recorded from completely unadducted larynges (N = 10, *Protocol 1*), calculated the mean Shannon’s entropy during maximum flow and used this value (0.7) for all other rat ramps to detect USVs. For mice, a Shannon’s entropy limit of 0.8 was used (based on visual inspection). Any continuous period of sound below these threshold levels, allowing for breaks of 1 time slice, were considered USVs. For time bins with USVs we extracted the peak frequency (*f*_*p*_) using the *tfridge* function.

### Comparison between model predictions of jet length and laryngeal geometry

#### Laryngeal geometry reconstruction and quantification

The diceCT scans were analyzed in Amira (Amira 5.2.1, 2009, Konrad-Zuse-Zentrum Berlin (ZIB), Visage Imaging Inc.). An oblique slice was placed in the sagittal plane, and another oblique slice was placed perpendicular to the first one, and overlapping the glottal opening (Fig. 1A). The slices were exported as TIF-images and imported into ImageJ (Version 1.52a, Wayne Rasband, National Institute of Health, USA) for measuring the following laryngeal dimensions: On the cross-section overlapping the glottal opening, we measured the glottal area, *A*_*gl*_, as the area of a polygon manually fit into the glottal constriction on the corresponding cross section (Fig. 3A, right). On the midline cross section, we measured the shortest and longest distances between the point of jet formation and the ventral intralaryngeal wall, *x*_*alar*_ and *x*_*max*_ (Fig. 3A, left), i.e. the range of jet lengths that could possibly fit in the larynx. Here we also measure the length of the opening of the ventral pouch, *x*_*cav*_. The point of jet formation was approximated as the point in the middle of the distance between the adducted arytenoids and adducted vocal folds (Fig. 3A, left). We assumed bilateral/axial symmetry for the jet, i.e. that its direction was parallel to the sagittal plane.

#### Mode analysis

In order to compare the jet length predictions based on the aerodynamic models corresponded with internal laryngeal geometry during USV production, we needed to identify which mode was extracted from the fUSVs. Both the jet impingement model and the alar edge-tone model predict the frequencies of several modes and therefore it was paramount to identify the mode numbers of fUSVs. We manually selected fUSVs where multiple modes were visible and compared the frequencies of other modes to the dominant frequency, *f*_*p*_, over time (Fig. 3B) using the tfridge MATLAB method on the spectrogram. The frequencies of the modes above the first one, *f*_*0*_, are given as *f*_n_=n ⋅ *f*_0_ (where n = 2, 3, 4, …). The difference in frequency of two adjacent modes is thus equal to *f*_*0*_ and the mode of the dominant frequency can be calculated as 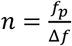, where Δ*f* is the difference between the frequency of the dominant mode and the closest mode, equal to *f*_*0*_ if the modes are of adjacent mode number. The frequency of the first mode was then calculated as 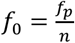.

#### Jet geometry predictions

The models predict different jet lengths:

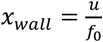, for the wall impingement model, (25)

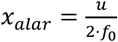, for the alar edge-tone model, (27)

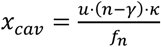, for the cavity-tone model, (35)

where *f*_*0*_ is the fundamental frequency, *u* is the mean convection speed of downstream moving coherent structures, approximated as the jet exit speed 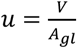(Fig. 3B, right), *V* is volumetric flow rate and *A*_*gl*_ is glottal constriction area.

Jet angle was determined by first fitting the predicted jet length between jet exit midpoint and the ventral intralaryngeal wall on the midline cross section. We then measured the angle between the resulting line and the midline of the cartilaginous glottis in ImageJ (Fig. 3A). As the jet length predictions for the edge-tone model were too short to reach the alar edge, we were unable to measure jet angle resulting from fitting *x*_*alar*_ between the jet exit midpoint and a point on the alar edge, but in theory, the alar edge-tone model predicts jet angles similar to *α*_*min*_ (Fig. 3A, left). For the cavity tone model, we did not investigate jet angle, as the model does not rely on jet formation.

### In vivo threshold flow estimate

We estimated tracheal air flow (*V*) during USV production in rats based on *in vivo* data. During quiet respiration *V* is 15-20 ml/s (64). However during USV production *V* reduces, which is seen in measurements of tracheal mass flow (30, 32, 52). We approximated *V* to be below 4 ml/s during USV production (Fig. 3A in (52)) for a 250-300 gram animal. We then linearly corrected for body size, which suggested that *V* during USV production for the animals used in this study was below 10 ml/s.

### Computational Fluid Dynamic simulations

We performed CFD (Computational Fluid Dynamic) simulations of air flowing through 3D-reconstruction of intra-laryngeal rat airways with and without a sphere digitally added to the ventral pouch (Fig. 3E-G & Fig. 4E-F). From the µCT scan of one of the larynges, the laryngeal airway was labeled in Amira. Under the experimental subglottal pressure condition, the mean jet speed is estimated to be about 40 m/s. The according Mach number (defined as Ma=u/c, where u is the mean jet speed, c=346 m/s is the speed of sound at 25°C) would be about 0.12. Therefore, the flow is modeled as an incompressible flow. The governing equations are the three-dimensional, unsteady, viscous, incompressible Naiver-Stokes equation as below

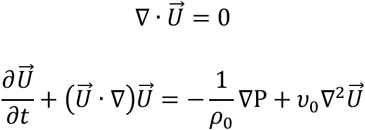

where 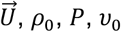 are the incompressible flow velocity, density, pressure and kinematic viscosity, respectively. v_0_=1.562×10^−5^ m^2^/s and *ρ*_*o*_=1.184kg/m^3^ at 25C. The computational solver employs the sharp-interface immersed-boundary method (37). The laryngeal wall is represented by triangular elements exported from Amira and smoothed through Meshlab. The governing equation is solved on non-uniform Cartesian grids, with finest grids at the glottal jet region. A 1.0 kPa subglottal pressure is applied at the subglottal entrance. A non-penetration non-slip wall boundary condition is applied at the laryngeal wall. Sensitivity studies on domain and grid size showed that the numerical solution converges with a minimum grid of 10 µm with a domain size of 26 x 32 x 28 mm with a 0.8% difference in the jet speed.

To test if the proposed flow scenario for the alar edge model is physically plausible, we performed CFD simulations of air flowing through a previously published intra-laryngeal rat airway (27). We obtained the 3D geometry from Morphobank (www.morphobank.org, project ID 2686, Morphobank media number M451228) and smoothed it in Meshlab. Simulation conditions were as listed above. We applied a 0.9 kPa subglottal pressure at the subglottal entrance.

### Statistics

All statistical testing was performed in MATLAB (MATLAB 2018a, MathWorks, USA). A two-tailed t-test was performed to test if the slope of the frequency-to-flow relationship differed before and after fixation of the larynges. As the core of the experiment was to predict jet length from peak frequency, mode number, and jet speed, we compared *x*_*wall*_ predictions between fixed and fresh larynges. For two of the fresh larynges, the mode number was difficult to determine and we decided to perform the tests with mode numbers that fell within the range we saw for the fixed larynges and that gave the predictions that best matched the corresponding predictions in the fixed larynges. We performed a two tailed t-test comparing the difference between the *x*_*wall*_ predictions from the fixed and fresh larynges to zero.

To compare the wall impingement model and alar edge-tone models’ jet length predictions to each other and to the internal laryngeal geometry during USV production, we performed two-tailed paired sample t-tests to compare predicted and measured *x*_*wall*_, *x*_*alar*_, and *x*_*cav*_, respectively.

We compared the peak frequencies of the fUSVs produced with and without a sphere filling the ventral pouch and blocking the alar edge using two-tailed paired sample t-tests on mean, maximum, and minimum frequencies of the fUSVs produced in the two treatments and for the two sets of animals (rats and mice).

### Quantitative motor control model

We constructed a quantitative data-driven model to capture how the activity of respiratory muscles (RM, mainly the diaphragm muscle) and a combination of intrinsic laryngeal muscles affect the main control parameters of our aerodynamic model: jet speed (*u*) and jet length (*x*). Subglottal pressure increases linearly from 0 to 5 kPa with RM activity. Tracheal flow (*V*) was predicted as a function of glottal area, tracheal diameter (measured from dice-CT scans), and subglottal pressure by assuming the glottal constriction to constitute a tube with an obstruction (65). We compared this obstruction model a ramp for a fixed larynx where glottal area, flow and pressure were known. The model prediction aligned well with experimental data (Fig. 5C).

State-of-the-art measurements and 3D models of vocal fold adduction on canine larynges (48-50) show how shortening of the adductor and abductor muscles sets glottal area. Based on these insights, we modeled glottal area as sum of the membranous glottis (area between the vocal folds) and cartilaginous glottis (areas between the arytenoid). The area of the membranous glottis was set by Thyroarytenoid (TA) activity and the cartilaginous glottis is set by TA and a combination of posterior cricoarytenoid (PCA) and interarytenoid (IA) muscles:

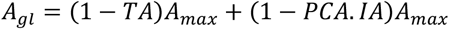

where *A*_*max*_ was measured from dice-CT scans (Fig. 3A). Because we lacked data on interaction between the TA and PCA.IA parameters we assumed them to be coupled.

The jet speed was defined as tracheal flow divided by glottal area. Contraction of the CT muscle rotates the thyroid wall away from the glottal opening (50), thereby increasing jet impingement length, *x* (Fig. 5E). TA weakly counteracts this rotational action of the CT by shortening the vocal folds (50), thereby decreasing the impingement length:

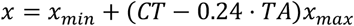

where *x*_*min*_ was defined as 50% of the minimum predicted impingement length and *x*_*max*_ 150% of the maximum predicted impingement length (see jet length prediction). The constant 0.24 was chosen as it results in an impingent length of zero at 100% TA activation and 0% CT activation.

By chaining these functions together, we can predict frequency as:

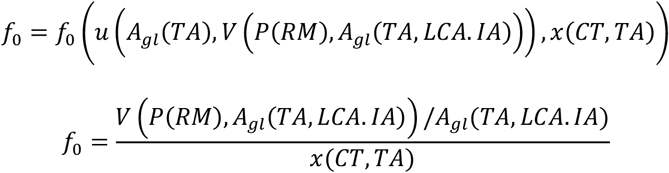

Lastly, we restricted the possible values for f0 by implementing the whistle stability criteria *d/x ≤ St < 1*, where *d* is jet diameter and *St = f0·d/u* is the Strouhal number (25). Simulations were implemented in Matlab and will be made available on Github. Because this is steady-state and not dynamical model the time representation in Fig. 5 is arbitrary and chosen to fit experimental data (44).

## Declarations

### Ethics approval

All experiments were conducted at the University of Southern Denmark and were in accordance with the Danish Animal Experiments Inspectorate (Glostrup, Denmark).

### Consent for publication

Not applicable.

### Availability of data and materials

The datasets used and/or analysed during the current study are available from the corresponding author on reasonable request.

### Competing interests

Authors declare that they have no competing interests.

### Funding

This research was supported by a Janet Waldron Doctoral Research Fellowship, the University of Maine (W.J.), and the Danish Research Council grant 7014-00270B (C.P.H.E.).

### Author Contributions

JH, AAG and CPHE designed research; JH, WJ, QX, XZ, MD, AAG and CPHE performed research; QX, XZ, MD, CPHE contributed new reagents/analytic tools; JH, WJ, QX, XZ, AAG and CPHE analyzed data; and JH, AAG and CPHE wrote the paper.

## Acknowledgments

We thank T. Christensen for technical support; M. Wöhr for discussion and comments on the manuscript.

## Additional Files

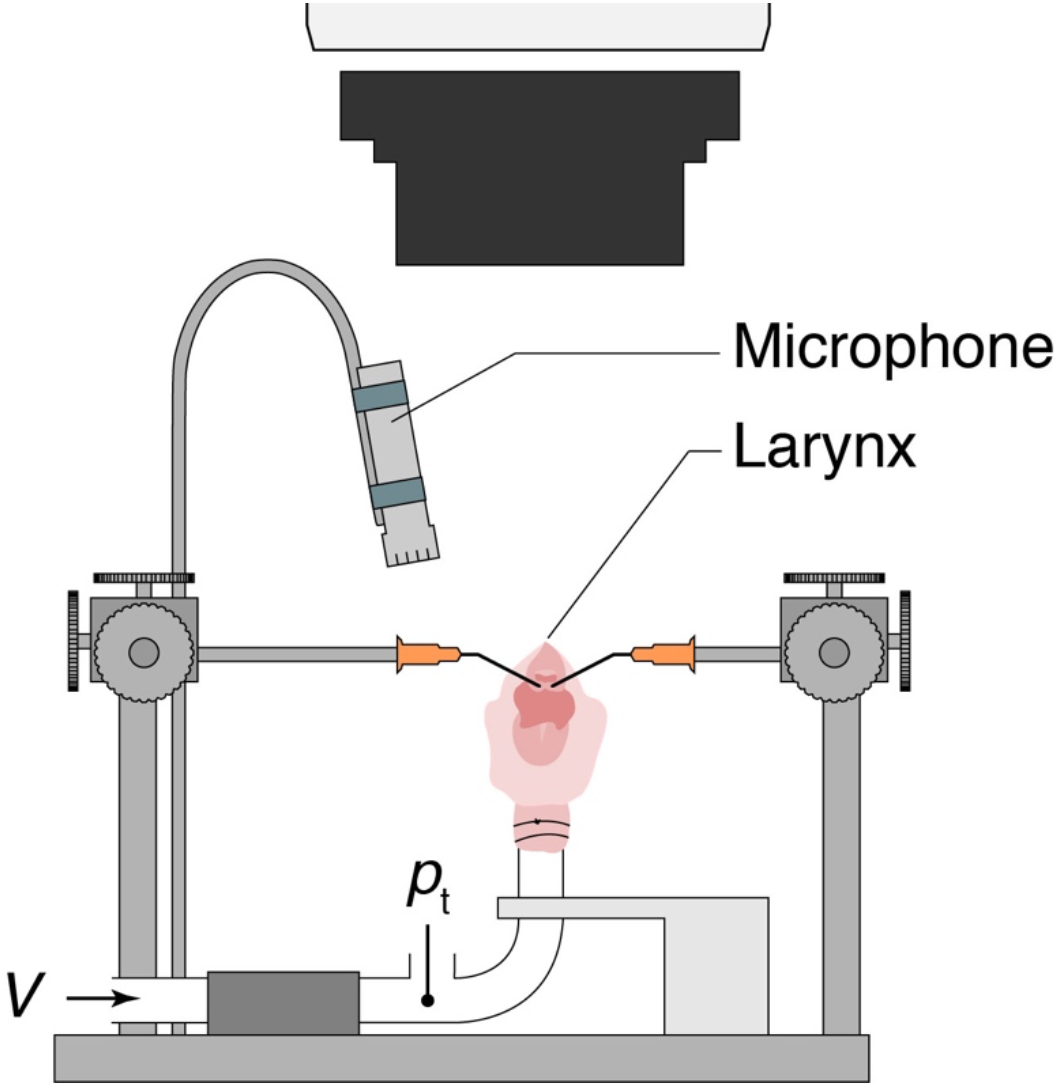

**Additional File 1**.

**File format: Figure S1.jpg**.

**Title: Schematic of in vitro larynx sound production setup**.

**Description:** The measurement position of tracheal pressure *p*_*t*_ and mass flow V are indicated. VF adduction is controlled with micro-manipulators.

**Additional File 2**.

**File format: Movie M1.jpg**.

**Title: CFD simulation of airflow through rat larynx with adducted vocal folds (Fig 3FG)**.

**Description**: Flow was simulated in a fixed 3D mesh of the laryngeal airway. This movie shows that a distinct jet is formed and impinges on the thyroid wall.

**Additional File 3**.

**File format: Movie M2.jpg**.

**Title: CFD simulation of airflow through rat larynx with filled ventral pouch (Fig 4EF)**.

**Description**: This movie shows that also with a filled ventral pouch, a jet forms that impinges on the thyroid wall with negligible effect on the jet length and angle.

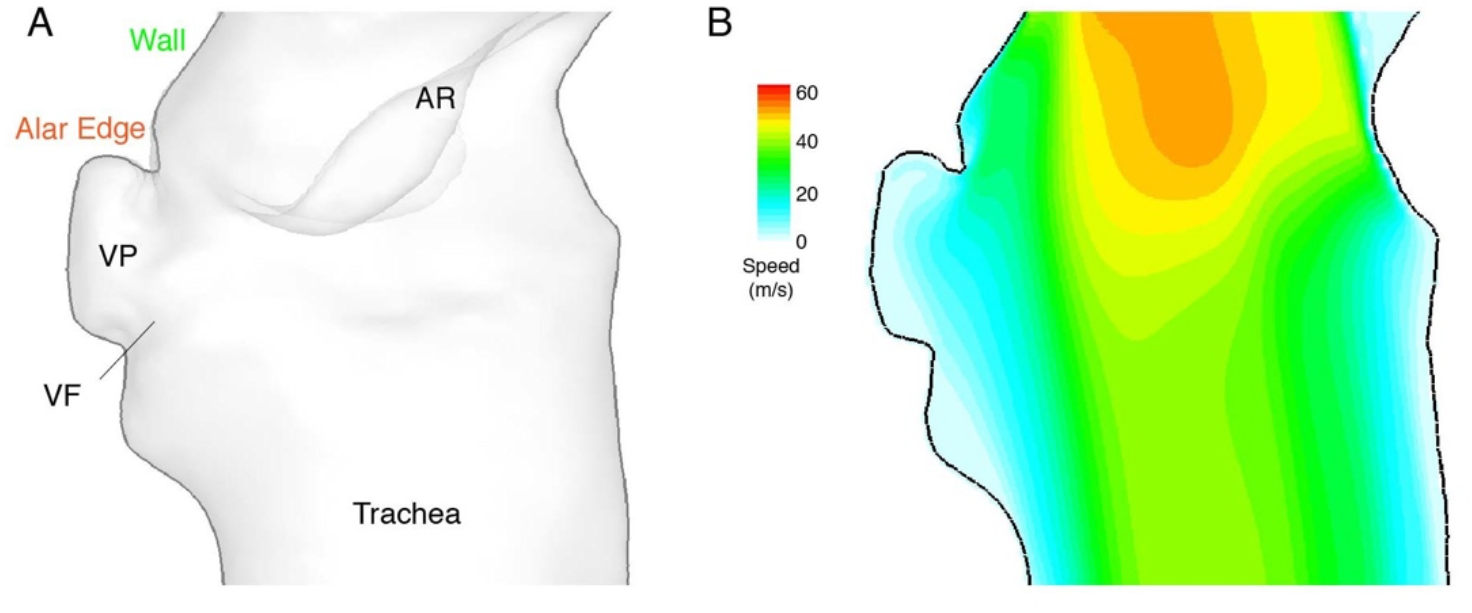

**Additional File 4**.

**File format: Figure S2.jpg**.

**Title:** CFD simulations does show no jet formation of a jet hitting alar edge or air circulation in ventral pouch in rat larynx with abducted vocal folds.

**Description: (**A) Geometry of previously published 3D airway reconstruction of the rat larynx (27). This air way geometry with abducted VFs was suggested to lead to USV production by either an air jet hitting the alar edge jet or by air circulation in the ventral pouch. (B) Modelled air speed through the air way (mass flow: 22 ml/s, tracheal pressure: 0.9 kPa) shows that no jet forms that hits either the alar edge or thyroid wall. Also, no air circulation takes place in the ventral pouch in this geometry. Therefore, the geometry of this air way does not support USV production following the alar edge model.

## Notes

### Competing Interest Statement

The authors have declared no competing interest.

